# Tight Turns of Outer Membrane Proteins: An Analysis of Sequence, Structure, and Hydrogen Bonding

**DOI:** 10.1101/304287

**Authors:** Meghan Whitney Franklin, Joanna S.G. Slusky

**Author notes:** Corresponding Author: Joanna S.G. Slusky.

## Abstract

As a structural class, tight turns can control molecular recognition, enzymatic activity, and nucleation of folding. They have been extensively characterized in soluble proteins but have not been characterized in outer membrane proteins (OMPs), where they also support critical functions. We clustered the 4-6 residue tight turns of 110 OMPs to characterize the phi/psi angles, sequence, and hydrogen bonding of these structures. We find significant differences between reports of soluble protein tight turns and OMP tight turns. Since OMP strands are less twisted than soluble strands they favor different turn structures types. Moreover, the membrane localization of OMPs yields different sequence hallmarks for their tight turns relative to soluble protein turns. We also characterize the differences in phi/psi angles, sequence, and hydrogen bonding between OMP extracellular loops and OMP periplasmic turns. As previously noted, the extracellular loops tend to be much longer than the periplasmic turns. We find that this difference in length is due to the broader distribution of lengths of the extracellular loops not a large difference in the median length. Extracellular loops also tend to have more charged residues as predicted by the charge-out rule. Finally, in all OMP tight turns, hydrogen bonding between the sidechain and backbone two to four residues away plays an important role. These bonds preferentially use an Asp, Asn, Ser or Thr residue in a beta or pro phi/psi conformation. We anticipate that this study will be applicable to future design and structure prediction of OMPs.

## Highlights

- The tight turns of OMPs differ structurally and sequentially from soluble proteins.
- Soluble and OMP turn differences are from constraints of the β-barrel and membrane.
- Different twists in OMP and soluble β-strands results in different turn types.
- OMP turns use D,N,S, and T residues in the B or P φ/ψ regions for hydrogen bonding.
- OMP tight turn hallmarks will assist the design of novel, functional OMPs.

## II. Introduction

Tight turns are often considered the third type of secondary structure after α-helices and β-sheets. Capping other structure types, tight turns are defined by their reversal of protein chain direction [1]. A quarter of all amino acids in proteins are engaged in this highly hydrogen-bonded structure type [2]. Tight turns are disproportionately represented at the surface of proteins where their high composition of charged and polar amino acids makes them frequent sites for molecular recognition [3]. Because of their biological activity and the stability imparted by their network of hydrogen bonds, many pharmaceuticals have been designed to mimic tight turn structure [4]. Tight turns also play an important role in folding and have been shown to nucleate folding of proteins [5] and nucleate the formation of beta hairpins [6].

Because of their importance in folding nucleation and their utility in molecular recognition, the tight turns of globular proteins have been well-characterized. With just 4 residues, β-turns are the most well-described of the tight turn lengths [7–11]. Five-residue tight turns (α-turns) and six-residue tight turns (π-turns) have also been well characterized [12–14].

Though the tight turns of outer membrane proteins have not yet been characterized, the utility of tight turns is no less prominent in outer membrane proteins than it is in soluble proteins. In the outer membrane, tight turns have been shown to control channel gating [15–17], receptor binding [18–20], outer membrane protein insertion [21] and possibly initiate outer membrane protein folding *in vitro* [22].

Outer membrane proteins share the antiparallel β-barrel fold. Because of the homogeneity of this topology, almost all connections between β-strands in the barrel fall into the category of tight turns with a small percentage making longer plug domains. In outer membrane β-barrels (OMBBs), tight turns on the extracellular side of the membrane are called ‘loops’ and the tight turns on the periplasmic side of the membrane are called ‘turns’ (Fig 1A, inset). We will refer to the general category of tight turns in OMBBs as ‘strand connectors’ to avoid confusion that tight turns might only refer to those connectors on the periplasmic side.

**Figure 1.**
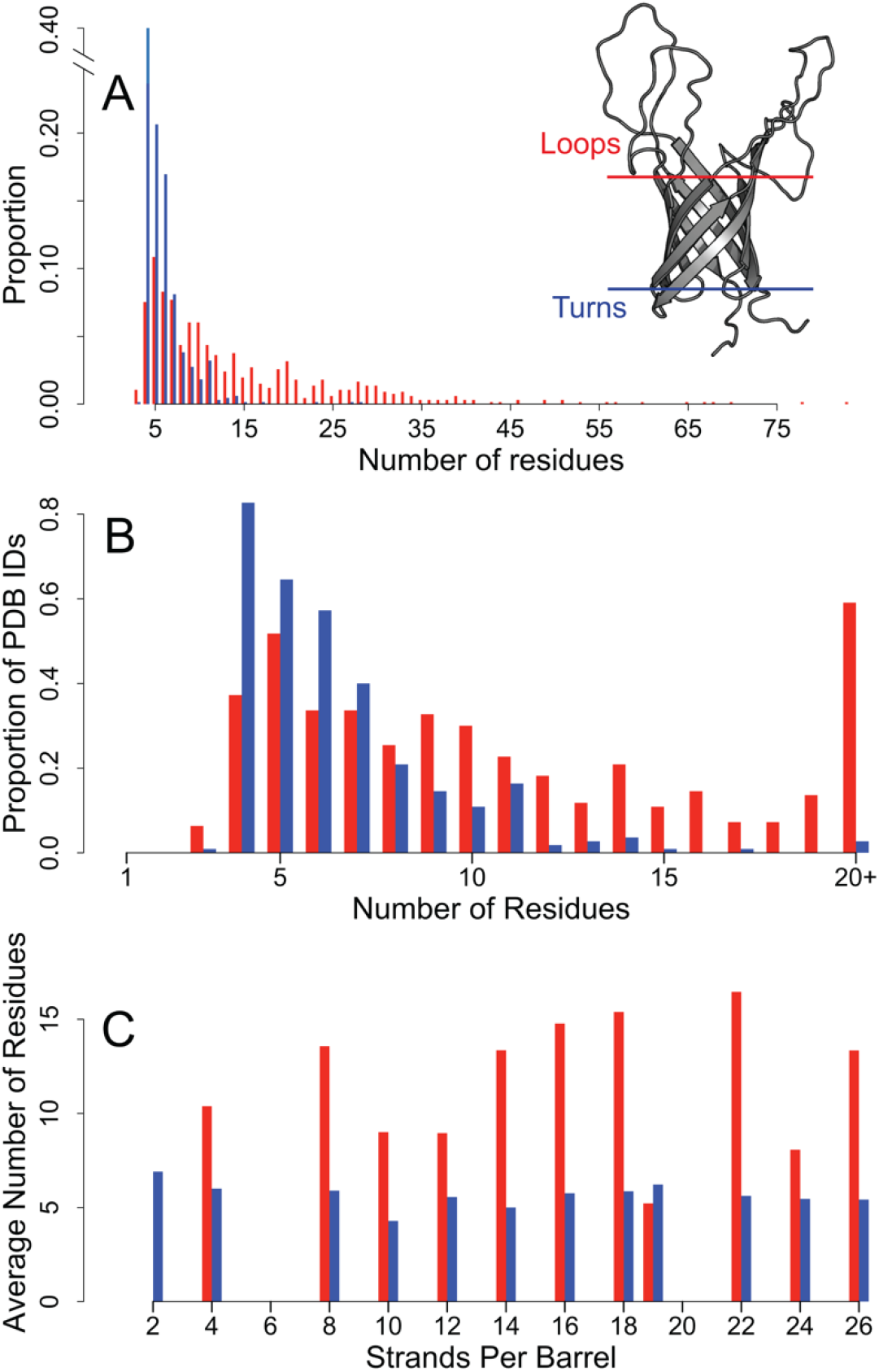
Comparison of distribution of loop lengths and turn lengths. Loop data are shown in red bars, turn data shown in blue bars. Distributions exclude excluding incomplete structures and turns with extensive secondary structure. **A)** Left: distribution of strand connector lengths. Right: The structure of OmpA (PDB: 2GE4) is shown with the membrane-bound region approximated by the blue and red lines. **B)** The proportion of PDBs in our dataset that contain a strand connector of each length. **C)** The average number of residues in the strand connectors grouped by the number of strands per chain per barrel.

In this study, we categorize the tight turns/strand connectors of OMBBs to determine what the distribution of structures are, what amino acids create those structures, and what types of hydrogen bonding impart stability to this fold. For strand connectors with lengths from 4-6 residues, we clustered them into groups based on the phi/psi angles of the non-terminal residues. We then describe these structural clusters with respect to structural composition, sequence composition, and hydrogen bonding preferences. We find different preferred structure types in the strand connectors of OMBBs than in the tight turns of soluble proteins. Moreover, we find different preferred amino acid compositions in these clusters. We anticipate that a thorough understanding of OMBB strand connectors will enable protein design to yield better folded and more functional OMBBs.

## III. Results

### A. Length-Specific Trends

From our outer membrane protein dataset of 110 proteins, we identified 779 extracellular loops and 688 periplasmic turns. We then rejected any incomplete strand connectors and turns that formed plugs or large periplasmic structures. 81% of the incomplete strand connectors were extracellular loops. As a result, this leaves 663 extracellular loops and 654 periplasmic turns.

A characteristic difference between periplasmic turns and extracellular loops is that the extracellular loops are longer [23]. This difference is so stark that one can visually determine the directional topology of an OMBB using this rule for almost all OMBB crystal structures. By graphing the strand connector lengths, we see that the mode of the two groups are similar while the distributions are different (Fig 1A). Turns have a mode of four amino acids and loops have a mode of 5 amino acids. More than half the loops contain 10+ residues while 96% of the turns are shorter than 10 residues. Like the overall lengths, the distribution of loops within PDBs is much wider than that of turns (Fig 1B); more than 80% of the PDBs in the data set contain a 4-residue turn while nearly 60% contain a loop with 20 or more residues. All barrels, regardless of the number of strands, have approximately the same average turn length (Fig 1C). In contrast, the loop lengths are less homogeneous with the longest loops observed in the 16-,18-, and 22-stranded barrels. For further categorization we refined the strand connectors to the groups of 4-6 residues. We find 510 turns (65% of all turns) of 4-6 residues and 177 loops (26% of all loops) of 4-6 residues.

To determine whether the sequence content of the loops differed from the turns, we permuted the assignment of each sequence as loop or turn and calculated the sum of the differences between amino acids at each position (see Methods). We find that when all positions are considered for each length, the amino acid content of the loops and turns are different (p ≈ 1×10^−4^, p ≈ 0, p ≈ 0 for the 4-, 5, and 6-residue strand connectors respectively).

#### 1. 4-residue strand connectors

By far the largest population of strand connectors are the 4-residue strand connectors. In our dataset, the 4-residue turns (n=264) and loops (n=50) are distributed into 5 clusters (Fig 2A). Cluster median angles and representative loops and turns are shown in Table S2. Of these five clusters of strand connectors, four of these belong to the nine canonical tight turn types for soluble proteins [11]. However, five of the canonical soluble strand connector types are never seen in our data. More than 80% of soluble protein four-residue tight turns are connected by type I’ or type II’ turns [10, 24]; however, just 58% of loops and 39% of turns in our data fall into these two types (Fig 3A).

**Figure 2.**
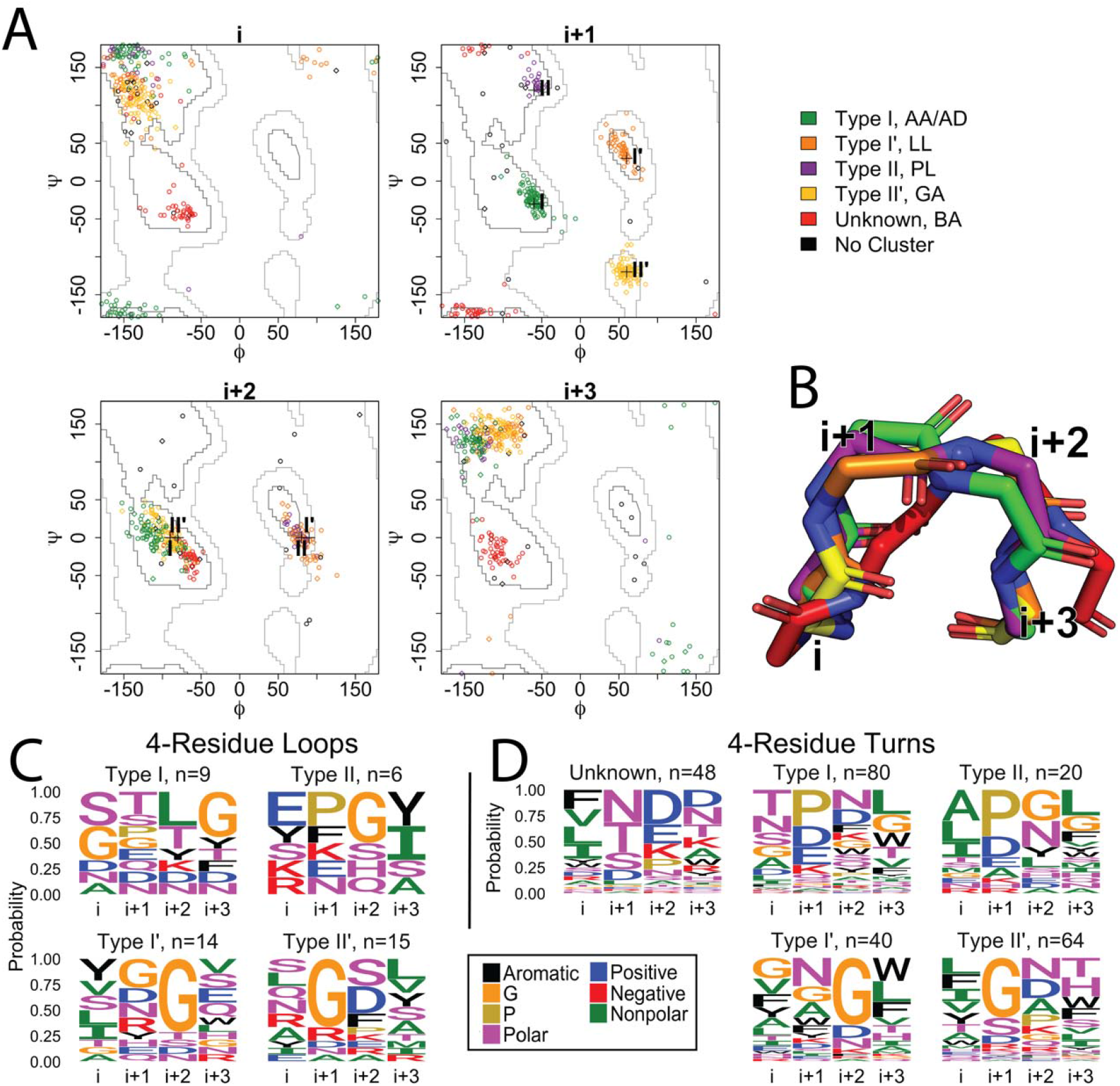
Four residue strand connectors. **A**) Clustering of the 4-residue loops and turns of OMBBs. Loops are shown with a diamond; turns are represented with circles. The plus symbols mark four of the previously defined ideal turn types I, II, I’, and II’ as noted by Guruprasad and Rajkumar (2000). Green: type I; orange: type I’; purple: type II; yellow: type II’; red: unknown; black: no cluster. **B**) Representative example of each cluster, colored as in A. The red cluster has terminal positions spaced more widely apart than the other four canonical tight turn types. **C and D**) Sequence logos for the 4-residue strand connectors. Each position is shown as a column; the number of sequences in a cluster is shown in the subtitle. Amino acid groupings are described in the methods. **C**) Sequence logos for the 4-residue loops. **D**) Sequence logos for the 4-residue turns.

**Figure 3.**
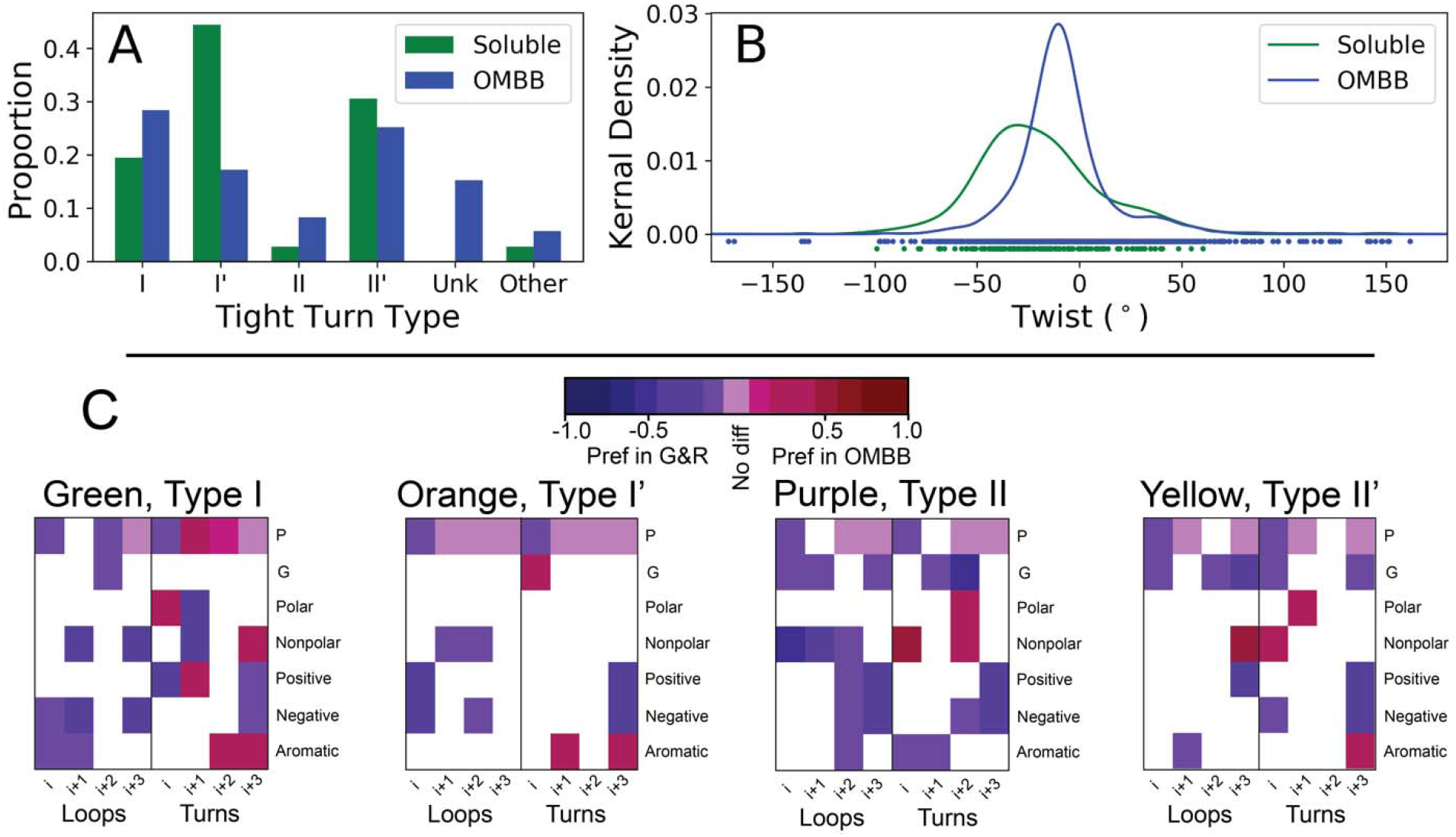
Comparisons between 4-residue OMBB strand connectors and 4-residue soluble protein tight turns. **A and B)** Comparison of the β-hairpins surveyed in Sibanda et al (1985) compared to the hairpins of OMBBs. **A)** Proportion of each of the 5 different types of 4-residue strand connectors observed in the OMBBs compared to those of the analogous 4-residue tight turns of soluble hairpins. “Other” refers to tight turns that do not fall into any of the other canonical β-turn types. **B)** Kernel density estimator of the twist of the hairpins connected by 4-residue tight turns in soluble proteins and in OMBBs. **C)** Statistically significant differences between Guruprasad and Rajkumar (2000, abbreviated G&R) and the OMBB clusters based on a permutation test. White blocks indicate no significant difference; pale purple blocks indicate a significant difference of zero between the proportion of of amino acid in the OMBB vs G&R. The remainder of the colored blocks indicate a statistically significant difference between the two. Red blocks have a higher propotion in OMBBs, indicating a preference in OMBBs, while blue blocks are have a higher proportion in G&R, indicating a preference in G&R. The amino acid groupings are defined in the methods.

In addition to the type I, II, I’, and II’ strand connectors, we identified a new cluster (red, Fig 2A) which was previously unknown. This novel strand connector type is only observed in the turns, comprising 18% of these structures. The terminal residues of this cluster of strand connector are 120 somewhat wider apart with an average distance of 7.8Å compared to the average distance of 121 the other 4-residue strand connectors at 5.3Å (Figure 2B).

Each cluster we found shows specific amino acid preferences at each position of the strand connector (Fig 2C, 2D). To determine whether the amino acid composition of each cluster was statistically different from the other clusters, we permuted the assignment of the sequences to the clusters (see Methods) separately within the loops and within the turns. We find that the amino acid composition of most of the 4-residue turn clusters are statistically different from each other while most of the 4-residue loop clusters are not statistically different from each other (Table S1).

It has been proposed that the high proportion of type I’ and II’ tight turns in soluble proteins is due to the complimentary twist of the strands [8, 10], resulting in type I’ and II’ being energetically favorable conformations and types I and II being energetically unfavorable conformations [25]. This is supported by our data where we find that the twist is less extreme for OMBB hairpins connected by 4-residue strand connectors than it is in soluble 4-residue tight turns (Fig 3B, p ≈0, see Methods).

We compared amino acid preferences in OMBBs to previous reports of positional amino acid preferences in each type of the tight turns of soluble proteins [26] (Fig 3C, Fig S1). The comparison demonstrates a change in composition of the terminal positions (i and i+3) which are defined as part of β-strands and which are often still in the membrane. Pro is rarely observed in terminal positions of OMBBs, although there is a strong preference for it in the central positions of the type I turns. The charged residues are also strongly dispreferred in the OMBBs at the i+3 position and occasionally the i position as well, while aromatic residues are often preferred in the i+3 position. The tendencies of the central residues are cluster-dependent. The tendencies of the central residues are cluster-dependent. Though the OMBB strand connectors are high in glycine they are less glycine rich than the soluble tight turns.

#### 2. 5-residue strand connectors

The 5-residue loops (n=72) and turns (n=135) fall into 10 different clusters (Fig 4A) in contrast to only five clusters for the 4-residue strand connectors. This doubling of clusters is due to the divergence of structural similarity between the loop and turn clusters in the five-residue strand connectors. We have named the clusters using the phi/psi notation previously devised [27] for the central positions (Fig 4A). While four of five 4-residue clusters are the same between turns and loops, only two 5-residue clusters are shared by turns and loops (AAL - blue and PLD - purple, Fig 4A). Cluster median angles and representative loops and turns are shown in Table S3. The largest turn cluster is the BAD cluster (orange, Fig 4A), which accounts for nearly 33% of the turns. The largest loop cluster is the PLD cluster (purple, Fig 4A) with 28% of the strand connectors. Amino acid content of most clusters is significantly different from the content of other clusters as determined by a permutation test (Table S1).

**Figure 4.**
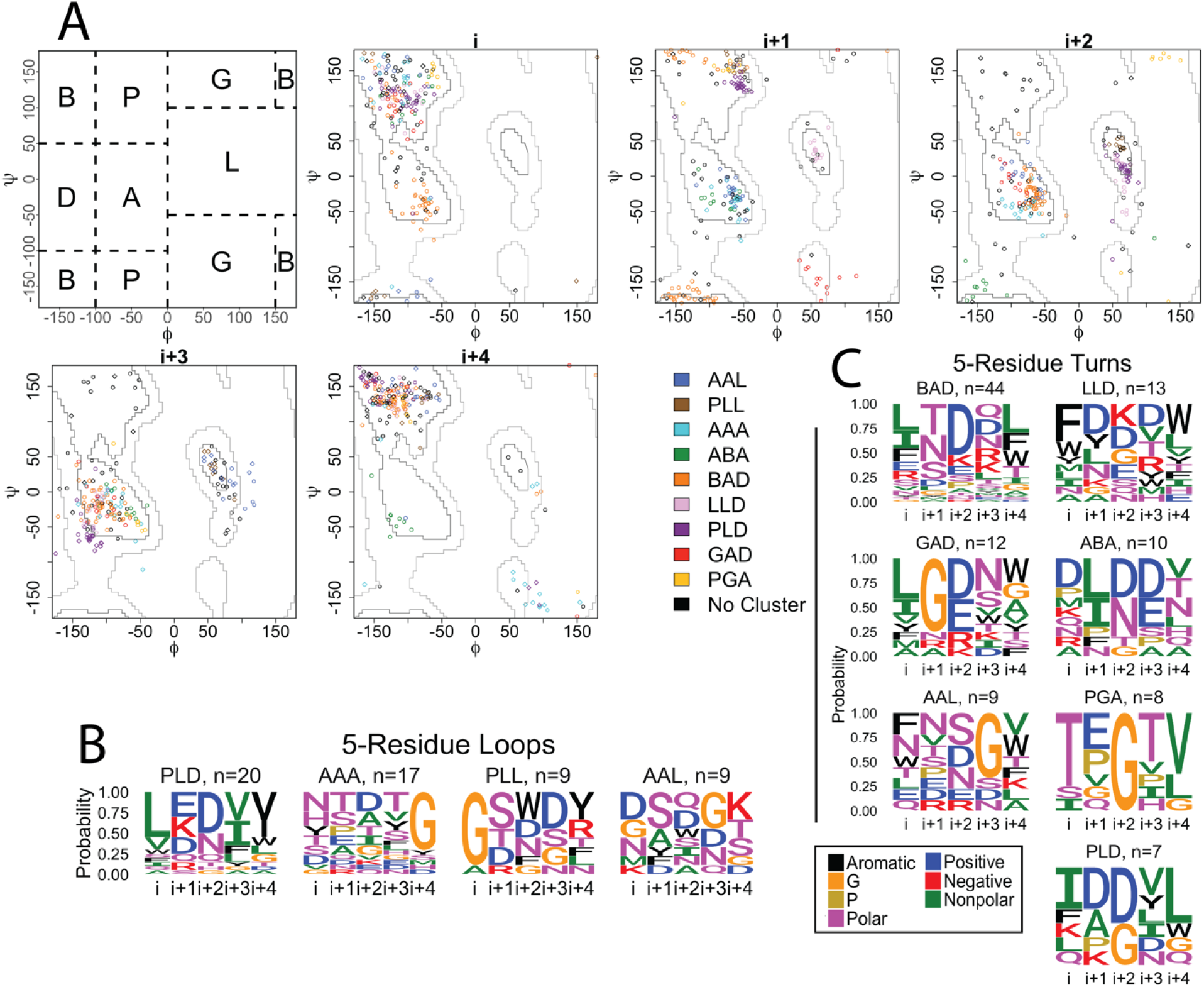
5-residue strand connectors. **A)** Clustering of the 5-residue loops and turns of OMBBs. Loops are shown with a diamond; turns are represented with circles. The first panel shows the division of the Ramachandran plot into regions according to North et al. 2011. The other 5 plots show the phi/psi angles for each residue in the strand connector. **B)** Sequence logos for the 5-residue loops. **C)** Sequence logos for the 5-residue turns. Sequence logos are colored and grouped as in Figure 2.

Unlike the 4-residue strand connectors, for which four of the five clusters have been previously documented, the clusters of 5-residue strand connectors of OMBBs show greater variation from previous reports of soluble tight turns. 15 clusters are described for soluble proteins covering 80% of the tight turns observed [28]. Of these, only four - AAA, AAL (called AAa), BAD (called BAA), and GAD (called pAA) - map to any of the 5-residue strand connector clusters we identify. These previously identified structures encompass just 38% of our structures. Therefore, it is difficult to calculate any sequence similarity between the OMBB strand connectors and the strand connectors of soluble proteins.

#### 3. 6-residue strand connectors

Like the 4- and 5-residue strand connectors, we find differences in sequence and structure between 6-residue loops (n=55) and 6-residue turns (n=111). Cluster median angles and representative loops and turns are shown in Table S4. Of the 7 clusters we identified, only one cluster (green, Fig 5A) is shared between the loops and the turns, accounting for 44% of the loops and 19% of the turns. The other 6-residue clusters contain only turns; the ADAD (blue, Fig 5A) cluster is the largest turn cluster with 21% of the structures. We find that the amino acid content within most clusters is statistically 178 different than the other clusters as determined by a permutation test (Table S2).

**Figure 5.**
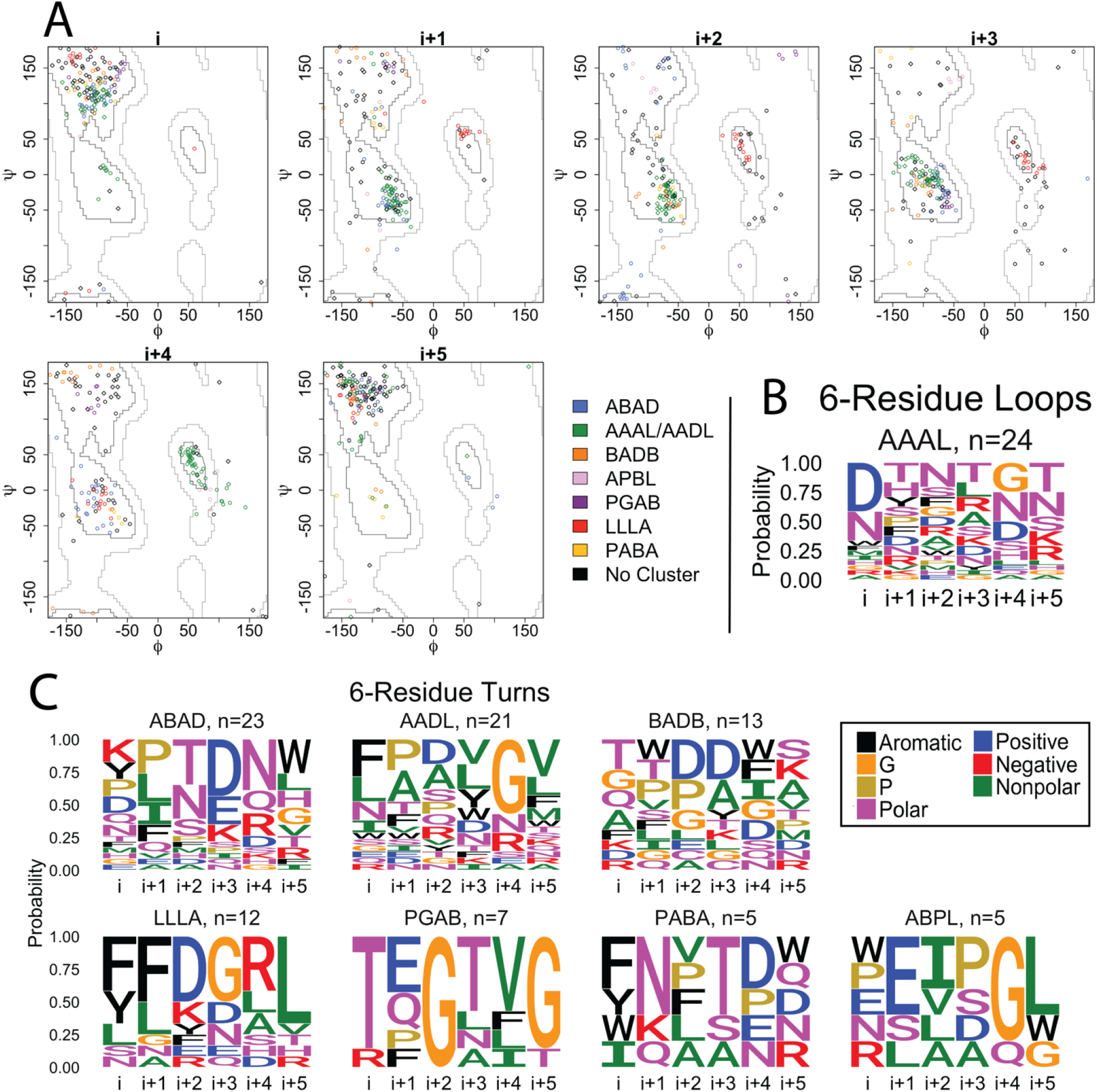
6-residue strand connectors. **A)** Clustering of the 6-residue loops and turns of OMBBs. Loops are shown with a diamond; turns are represented with circles. Because the i+3 residue of the green cluster straddles the boundary between the D and A regions, the green cluster is labeled AAAL in the loops and AADL in the turns. **B)** Sequence logos for the 6-residue loops. **C)** Sequence logos for the 6-residue turns. Sequence logos are colored and grouped as in Figure 2.

These 6-residue structural clusters show some similarity to previously described soluble tight turns [13] with 2 of the 9 reported soluble tight turn types appearing in our OMBB strand connector clusters. These two structure types comprise 30% of our total OMBB strand connectors. Specifically, the shared clusters are APBL (AEEa) and AAAL/ AADL(AAAa, green). However, the small number of counts with 30% overlap is not enough to compare sequence similarity between soluble and membrane proteins at any level of statistical significance.

### B. Backbone-backbone hydrogen bonding, terminal positions

Backbone-backbone hydrogen bonding is not a requirement to be considered a strand connector. However, since the loops connect two adjacent β-strands in a β-sheet, there are a high proportion of bonds between the last residue of the first strand (i) and the first residue of the last strand (i+n-1). The carbonyl oxygen of the first residue is often ideally placed to hydrogen-bond to the nitrogen of the terminal loop position and vice versa, although this preference is less pronounced in the longer strand connectors (Fig S2). The presence of this double or single backbone hydrogen-bond has been used to differentiate and categorize soluble tight turns [24].

In the 4-residue loops and turns, approximately 80%-90% of the structures in all of the clusters contain a double backbone hydrogen bond between the terminal residues. The notable exception is the previously unknown cluster which has almost no backbone-backbone bonding but instead participates in extensive sidechain-backbone hydrogen bonding (discussed below). The lack of terminal position backbone-backbone hydrogen bonding is likely because of the longer distance between these two amino acids. The 5-residue loops and turns generally only have a single hydrogen bond between the terminal residues, consistent with previously observed patterns [24], but this can vary widely from cluster to cluster. For example, nearly 90% of the structures in the LLD cluster (pink, Fig S2) and 80% of the PLD cluster (purple, Fig S2) have double hydrogen bonds, while the BAD (orange, Fig S2) and AAL (blue, Fig S2) clusters only contain a single hydrogen bond. In the six-residue strand connectors, approximately 80-90% of the LLLA turn cluster (red, Fig S2) and AADL loop (green, Fig S2) cluster have double hydrogen bonds. The equivalent green turn cluster AAAL (also green, Fig S2) also contains terminal backbone bonds, but these are more weighted towards single bonds. The remaining 6-residue turn clusters have almost no backbone:backbone hydrogen bonds between terminal residues.

### C. Backbone-backbone hydrogen bonding, non-terminal positions

In our strand connectors, backbone-backbone hydrogen bonding patterns also exist outside the terminal amino acids. In fact, these bonds are especially common in the 5- and 6-residue strand connectors, which show more variable patterns of bonding than the 4-residue strand connectors. In the 4-residue strand connectors more than a third (35% of turns and 42% of loops) of the structures contain a bond between the carbonyl oxygen of the i position and the backbone nitrogen of the i+2 residue (abbreviated i O:i+2 N); this is most common in the type II’ and type II strand connectors. In the 5- and 6-residue strand connectors, the carbonyl oxygen of the first position most often hydrogen bonds to residues that are two to four positions away (i O:i+2 N through i+4 N). Like terminal bonding, these patterns can be heavily dependent on the particular cluster; for example, the 5-residue PLD (purple, Fig S2) and PLL (brown, Fig S2) clusters have extensive backbone bonding within the loop, while the 5-resisdue ABA (green, Fig S2) cluster has almost no backbone-backbone bonding.

### D. Side chain-backbone hydrogen bonding

Positions within clusters that have phi/psi angles in the P and B region (Fig 4A) dominate the side-chain-backbone hydrogen-bonding. These positions include the 4-residue previously unknown (red) i+1 position, 5-residue BAD (orange) i+1 position, and 6-residue PGAB (purple) in position (Fig 6C). In these positions, sidechain-backbone hydrogen bonding correlates to the probability of the presence of polar amino acids Asn, Thr, Ser, or negatively charged Asp (Fig 6A, R^2^ = 0.686). These side chains tend to hydrogen bond with the oxygen or nitrogen of backbones that are two to three positions towards the C-terminus from them (47% of sidechain-backbone hydrogen bonds; Fig 6C). In rare cases the side chains form hydrogen bonds with backbones that are 4 positions away from them (4.9% of sidechain-backbone hydrogen bonds) or that are towards the N-terminus (17.6% of sidechain-backbone hydrogen bonds). A notable exception to the correlation between side chain-backbone hydrogen bonding and the proportion of NTSD residues is the i+1 position of the PLL (brown) loop cluster. This position, which falls into the P region, contains 89% N, T, S or D and yet shows no sidechain-backbone hydrogen bonding. Upon inspection of the structure the reason is clear - the sidechain of the i+1 position points outwards rather than towards the middle of the loop, preventing the possibility of any hydrogen bond forming.

**Figure 6.**
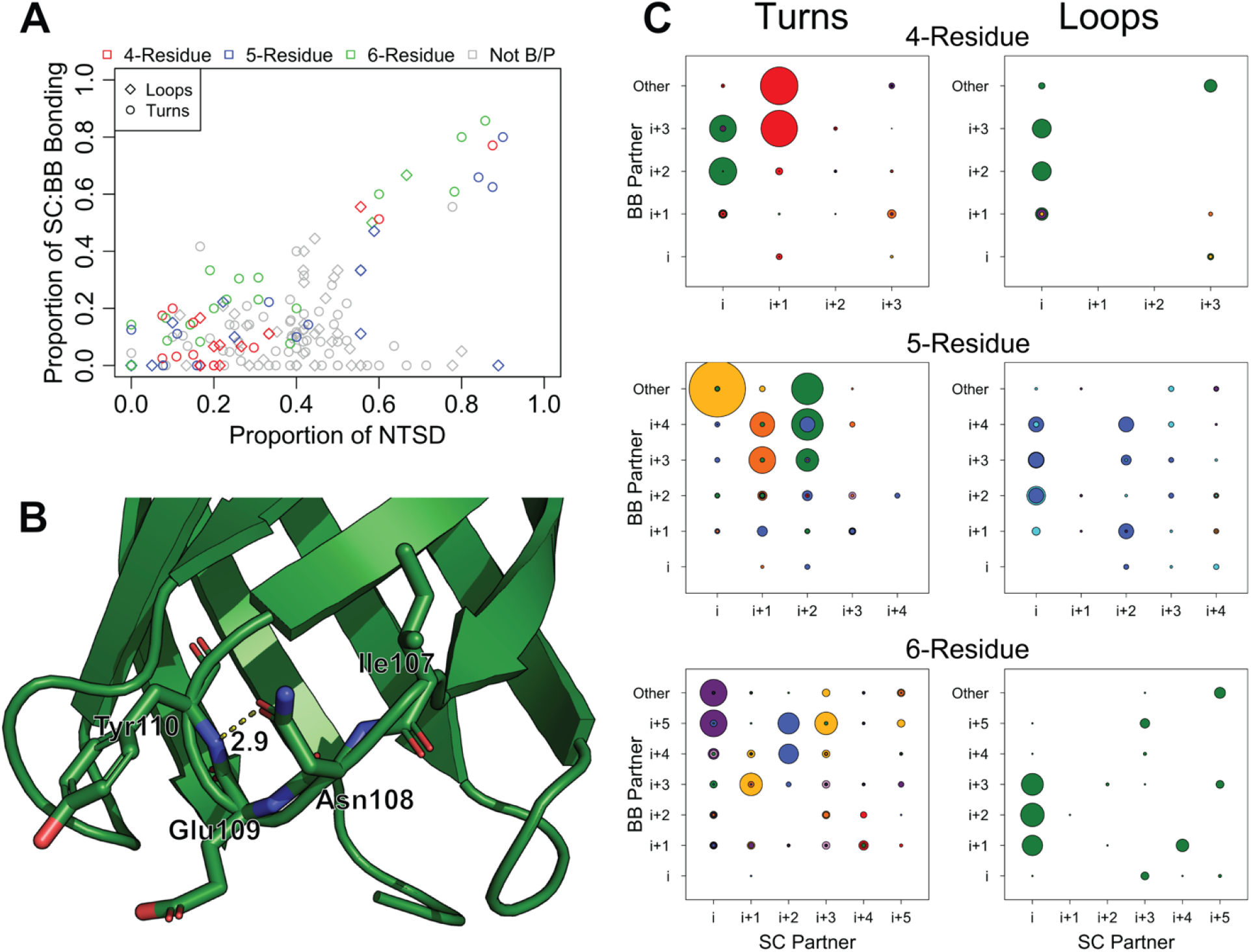
Side chain backbone bonding in strand connectors. **A)** Proportion of Asn, Thr, Ser, and Asp (NTSD) in any position compared to the proportion of sidechains donated to a side-chain:backbone hydrogen bond at that position. Positions that fall outside the B or P region of the Ramachandran plot are shown in grey. **B)** Turn 2 of PDB ID 3qra. The side-chain:backbone hydrogen bond between the side chain of i+1 Asn108 and the backbone of i+3 Tyr110 is shown in dotted lines. Glu109 is the i+2 residue and its side chain points directly outward. **C)** The proportion of sidechain:backbone hydrogen bonding at each pair of positions. The identity of the sidechain partner is shown along the x-axis; the identity of the backbone partner is shown along the y-axis. “Other” indicates hydrogen bonding to a backbone partner outside the loop. The radius of the circles is equivalent to the proportion of structures at that position involved in bonding. Circles are colored to match the clusters marked on the Ramachandran plots in Figs 2, 4 and 5.

### E. Use of the right-handed side of the Ramachandran plot

Although 58% of our structures use the right-hand (positive phi) side of the Ramachandran plot, very few amino acids can access these regions. The right-hand side is divided into the G and L regions as previously shown (Fig 4A). Just Asp, Asn and Gly are capable of occupying these regions [29]. We find that strand connector positions that occupy the G or L region are enriched in Gly, Asp, and Asn (Fig 7). Positions that fall into the G region contain more than 70% Gly residues (Fig 7, left bars). The L region is more equally distributed between these three amino acids (Fig 7, right bars), containing approximately 1/3 Gly residues in both the loops and turns and nearly 14% Asn. However, the loops contain a higher proportion of Asp than the turns likely because of the charge out rule [30].

**Figure 7.**
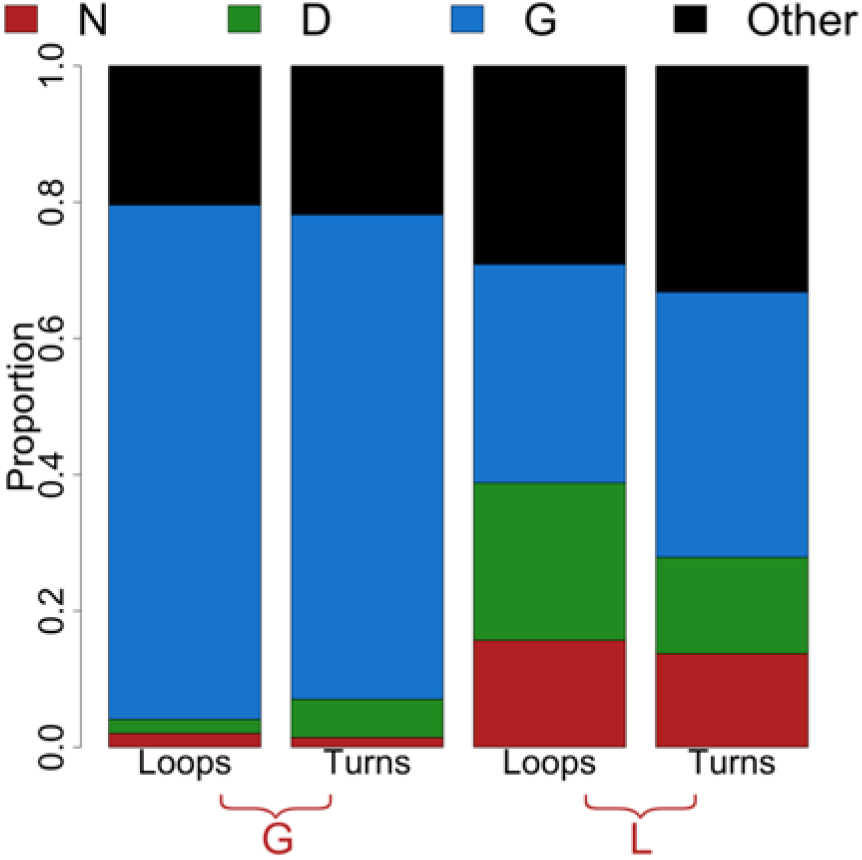
Proportion of Asp, Asn, and Gly in positions that fall into the G or L region of the Ramachandran plot. The remaining 17 amino acids are grouped into “Other”.

### F. Amino acid preferences at terminal positions

Specific amino acid preferences are found at the terminal positions. Because the terminal positions are frequently located in the membrane, both the loops and turns show an enrichment of nonpolar and aromatic residues (Fig 8). This trend is exaggerated in the terminal positions of the turns, which contains 55% of these amino acids while the loops only have 1/3 nonpolar and aromatic residues. The terminal positions also contain a decreased proportion of positive or negative amino acids; terminal positions have ∼10% charged residues, while the nonterminal positions contain nearly three times the charged residues. Finally, there is an absence of proline residues due to the β-strand nature of these positions. All three of these features reflect these positions’ secondary structure and their frequent localization in the membrane.

**Figure 8.**
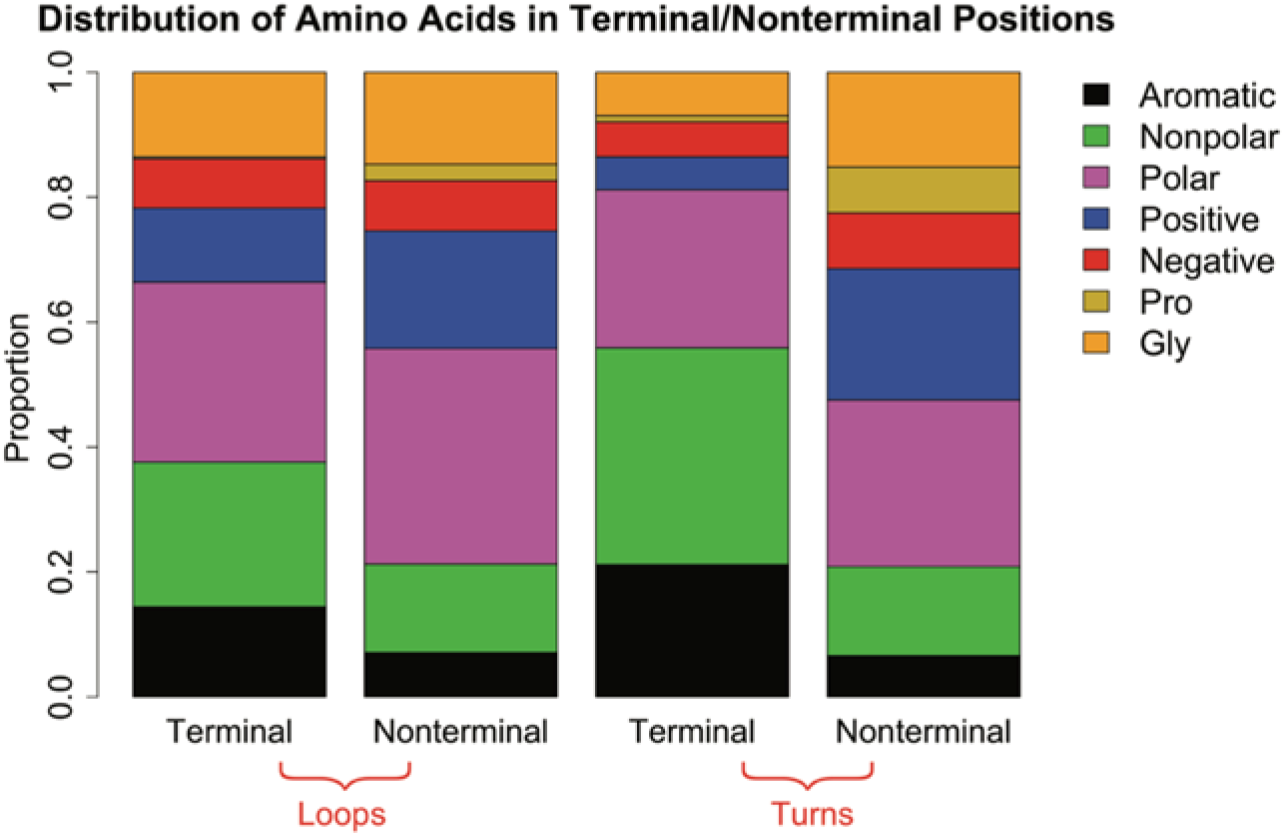
Proportion of residues that fall into each category in the terminal or nonterminal positions.

## IV. Discussion

Here we document structural, sequence and hydrogen bonding trends of strand connectors in OMBBs and compare those trends to the trends of tight turns in soluble proteins. We find preferences in structure, sequence, and hydrogen bonding for loops and turns of strand connectors in OMBB. We have compared loops to turns and structures from soluble proteins with structures from OMBBs to better understand the characteristics of the strand connectors in OMBBs.

In OMBBs we find that extracellular loops have a much broader distribution of lengths than the periplasmic turns. The mode length of a periplasmic turn is 4 amino acids and the mode length of an extracellular loop is 5 amino acids. However, because of the broad distribution of the loops we see much better representation of turns in the 4-, 5- and 6-residue length and therefore see more turn cluster types for each structure length.

Upon clustering the 4-, 5- and 6-residue strand connectors, we find that the longer the strand connector the more difficult it is to identify clusters. Of the 4-residue strand connectors, just 6% were outliers and did not fall into a cluster, while this proportion grows quickly to 24% outliers in the 5-residue strand connectors and 34% outliers in the 6-residue strand connectors. Lengths of 7 or more residues do not have sufficiently large populations to identify clusters. This is likely related to the greater number of permutations available. With 6 regions of Ramachandran space (Fig 4A), a four-residue strand connector has 6^4^ possible combinations, a six-residue strand connector has 6^6^ combinations, although not all of these combinations would be sterically possible.

### A. Comparison of OMBB strand connectors to soluble protein tight turns

#### 1. Structural differences

Structurally, outer membrane β-barrels put unique restrictions on strand connectors. The strands of OMBBs almost always lie 40° to the barrel axis [31] tilting rightward while the next strand is positioned leftward. This structural uniformity limits and changes the distribution of sequences and structures used to form connections between strands.

In comparing 4-residue soluble tight turns and OMBB strand connectors we find that there is a different distribution of structure type. Specifically, soluble tight turns most commonly contain the type I’ and type II’ tight turns [10, 24] followed by type I tight turns [26]. In contrast, we find that in OMBBs strand connectors contain 20-30% each of the type I, type II’ and our newly described red cluster, typed BA based on its Ramachandran regions (Fig 4A). The difference in turn type distribution is likely a result of the difference in twist distribution. Extreme twist favors the type I’ and type II’ turn types. OMBBs have a more homogeneous and less extreme twist which allows the more even distribution of tight turns we observe in the OMBBs.

Although we find differences between 4-residue strand connectors in soluble and OMBB proteins, the bigger differences are found in the bigger strand connector structures where we observe almost completely different sets of structures compared to the soluble proteins [13, 28]. Even in a survey specifically of soluble β-hairpins, half of the 5-residue turns are defined as ADL [24], which isn’t a cluster we observe in the OMBB strand connectors.

#### 2. Sequence differences

The amino acid preferences in terminal positions of OMBB strand connectors differ from terminal positions in soluble protein tight turns. The origin of the divergence of amino acid preferences is the localization of these terminal positions in the membrane of the OMBB strand connectors. These positions are defined as the last residue of the first strand and the first residue of the next strand. In soluble proteins, nonpolar and aromatic residues are not preferred in terminal positions and polar and charged residues are preferred. Due to the membrane-localized nature of the terminal positions, we find the opposite trend holds for OMBBs – nonpolar and aromatic residues are heavily favored while charged residues are much less preferred. Nonpolar amino acids are preferred in the membrane [32] and aromatic residues have previously been shown to prefer the headgroup region of the outer membrane [33].

### B. Hydrogen bonding

The presence of hydrogen bonds in the terminal residues of soluble tight turns has also been implicated increasing the rate of folding in soluble proteins [5] and we similarly find these bonds in OMBB strand connectors. OMBB strand connectors are capable of forming multiple hydrogen bonds, utilizing either the backbone or the sidechain hydrogen bond donors and acceptors. Side chain interactions of soluble tight turns have also been strongly implicated in the nucleation of beta hairpins [6]. We anticipate that the formation of tight turn hydrogen bonds might nucleate the bolding of OMBB hairpins as well.

We find side chain-backbone hydrogen bonding that is dominated by the amino acids NTSD in the B and P regions of the Ramachandran plot.

## V. Conclusion

In OMBBs strand connectors are involved in molecular recognition, channel gating, and membrane insertion. Here we find differences in sequence, structure, and hydrogen bonding between OMBB strand connectors and previously characterized soluble tight turns. We also compared periplasmic turns and extracellular loops of OMBBs and find differences between these two groups.

Most of the differences between OMBB strand connectors and soluble tight turns originate from the sequence constraints imposed by the membrane or the structure constraints imposed by the beta barrel fold. OMBB beta-strands are less twisted than soluble beta-strands which correlates with a difference in the distribution of previously described tight turn types that they can access. We find differences in sequence that are affected by localization in or near the membrane, e.g. more membrane proximal positions have a higher proportion for non-polar and aromatic amino acids. We also find sequence differences that are rooted in the structural differences because different backbone angles favor different amino acid identities. Finally, the hydrogen bonding preferences that we find are a product of both the sequence and structural components—backbone-backbone hydrogen bonding is dependent on structure type, but side chain to backbone hydrogen bonding is favored by Asn, Asp, Thr, and Ser residues in the beta or pro phi/psi regions.

The differences between OMBB loops and turns are more subtle. Many are linked to the different distribution of sizes of these structures though the underlying reason for this difference in size remains unclear.

We anticipate that this data could next be used to benchmark the various loop building and loop prediction software options for better application to outer membrane proteins. Function in these loops could be correlated to maintenance or divergence from the common structures and sequences. Ultimately, we hope the delineation of outer membrane protein tight turn hallmarks will lead to the design outer membrane proteins with novel functions.

## VI. Methods

### A. Loop Definitions

Our set of loops was extracted from the previously described set of 110 OMBBs at <50% sequence identity [31]. All amino acids were categorized by their phi/psi values. β-sheet amino acids were defined to have psi angles between −100° and 50° and the phi angles less than −100° for single-chain barrels and < −90° for multi-chain barrels. We also included amino acids with phi angles >0° and psi angles >150° as β-sheet amino acids.

To determine the strands, a combination of hydrogen bonding and a pattern of β-sheet residues was used. Because hydrogen bonds are often observed in loops of various lengths, DSSP definitions [34] were checked at the ends of each possible strand to ensure we hadn’t over- or under-extended the strand; residues defined as hydrogen-bonded turns (type “T”) were removed from strands while residues defined as extended strand (type “E”) were added. Following the convention for tight turns connecting soluble β-strands, loops were defined as the residues between the last residue of one strand and the first residue of the next strand, inclusively [10], regardless of the presence of any secondary structures. Because we are surveying OMBBs, we will refer to periplasmic tight turns as turns and extracellular tight turns as loops. Each OMBB contributes n/2 loops and (n/2)-1 turns where n is the number of strands in the barrel. This results in 779 loops and 688 turns for a total of 1458 strand connectors. Incomplete strand connectors and turns that formed plugs or large periplasmic structures were removed leaving 663 extracellular loops and 654 periplasmic turns.

The twist of strands was calculated using the definition of Murzin et al, 1994 [35], in which the twist is the dihedral angle between two residues in adjacent strands connected by a backbone hydrogen bond. The vectors are formed by four residues on two strands—two on each strand. On the first strand, we define the first residue. The second residue is the residue preceding the first residue on the peptide chain. On the second strand we define the residue hydrogen-bound to the first residue on the neighboring strand as the third residue. The fourth reside is the next residue on the peptide strand from the third residue. Together, the four residues form a U-shape. Within an OMBB, the neighboring strand can either be defined as the previous strand or the next strand. This results in two twist angles for each pair of residues – a forwards-looking and reverse-looking angle. Since the kernel density estimates of the distribution are similar, the data shown here uses the previous strand as the neighboring strand.

### B. Strand Connector Analysis

Strand connectors with missing residues in the structure file were designated as incomplete and excluded from further analysis. There are 143 incomplete strand connectors; just 27 of these are turns. We also excluded turns with more than 20 amino acids that contained more than two contiguous regions of 5+ residues classified as E, H or G by DSSP. This removed intra-barrel plug domains and the long periplasmic helices found in the efflux pumps such as TolC. Strand connectors were then sorted by length and the strand connectors of 4-6 residues were analyzed further. We used the sum of the chord distances between the phi/psi angles of the central, non-terminal residues to cluster each strand connector length in a similar fashion to North, et al. in their survey of antibody loops [27]. Residues are labeled from i to i+(n-1), where n is the number of residues of the strand connector.

For each turn/loop length, the strand connectors were clustered using HDBSCAN [36] and DBSCAN in the scikit-learn package for Python [37]. Conventionally, the terminal residues are excluded from clustering and describing the groups of any length. Values of eps (a minimum cluster density parameter) for DBSCAN were explored for each length and type. These two algorithms were chosen because they do not force all points into a cluster but exclude outliers as necessary. The optimum clustering algorithm was selected based on the silhouette score [38], Calinski score [39], and visual inspection of the clusters and number of outliers. The silhouette and Calinski scores were used because they require no knowledge of the “true” clusters. 4-residue clusters were named according to the 9 canonical turn types [11]. 5- and 6-residue clusters were named according to the regions in which the non-central phi/psi angles of the cluster centroid lie, following the delineation of North, et al (2011) (Fig 4A). The cluster assignments and phi/psi values for each sequence of 4-6 residues are in the supplemental file ClusterAssignments4_6Res.xlsx; the individual amino acid proportions by cluster are in the supplemental file AACompositionByCluster.xlsx.

### C. Amino acid groupings

Aromatic residues were defined as Phe, Trp and Tyr; nonpolar residues are defined as Ala, Ile, Leu, Met, and Val; polar residues are defined as Asn, Cys, His, Gln, Ser, and Thr; negative residues are defined as Arg and Lys; positive residues are defined as Asp and Glu. Proline and glycine are always considered separately because of their unique ability to access different parts of the Ramachandran plot.

### D. Comparison of different groups

Permutation tests with n=10,000 were performed by permuting group assignments of sequences to determine whether two groups of strand connectors were different. For each pair

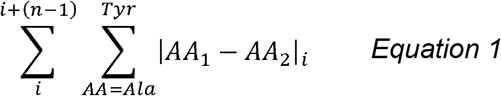

of groups, we calculated the sum of the differences between each amino acid at each position (Eq 1), where AA_1_ and AA_2_ represent the proportion of the same amino acid in two groups, i is the first position within the strand connector, and n is length of the loop. Permutation tests were performed to determine whether the amino acid composition of loops and turns of each length were statistically different (p<= 0.05) and to determine if the individual amino acid composition of clusters of a length and type were different from each other. For the differences between clusters, p-values <= 0.05 were considered statistically significant with a Holm-Bonferonni correction to minimize false positives.

To compare the loops to the turns, the assignment of strand connector type was permuted. For comparison of the Guruprasad and Rajkumar (2000) data, the amino acids within each position was permuted rather than the sequences and the sum differences were calculated as above. To determine whether the clusters within a strand connector type and length differed, the assignment of structures to a cluster were permuted. Finally, to determine if the distribution of twists in the soluble beta-hairpins differs from the twist of OMBBs, we permuted the assignment of soluble or OMBB (n = 10,000) and calculated the shared area under the resulting histograms with a bin size of 10°.

### E. Determination of hydrogen bonding

For each atom within each residue of each strand connector, possible hydrogen bonding was considered for any atom within +− 3 residues of the loop, inclusive. Within this distance-cut off, a geometry-based definition was used to determine whether two atoms (atom1 and atom2) could be hydrogen-bonded. The definition relies on the definition of two ‘previous atoms’, one to each of the other two atoms of interest. For the backbone nitrogen, the previous atom was set to the C_α_; for the C_α_, the previous atom was set to the backbone nitrogen.

Three conditions are necessary for two atoms to be considered hydrogen-bonded. First, the angle formed by atom1, atom2, and the previous atom to which atom2 was bonded (with atom2 at the center) is > 90°. Secondly, the angle formed by atom2, atom1, and the previous atom to which atom1 was bonded (with atom1 at the center) is also > 90°. Finally, the distance between atom1 and atom2 must be ≤ 4A. Hydrogen atoms and all carbon atoms (except the C_α_) were excluded from consideration.

## VII. Acknowledgements

Professor Eric Deeds for helpful discussions and members of the Slusky lab for helpful edits. NIH awards DP2GM128201, P20GM103418, P20GM113117, T32-GM008359, NSF MCB160205, The Gordon and Betty Moore Inventor Fellowship, KU-startup.

